# Shared and distinct genetic features in human and canine B-cell lymphomas

**DOI:** 10.1101/2021.10.14.464277

**Authors:** Krysta M Coyle, Tiana Hillman, Matthew Cheung, Bruno M. Grande, Kevin R. Bushell, Sarah E. Arthur, Miguel Alcaide, Nicole Thomas, Kostiantyn Dreval, Stephanie Wong, Krishanna Campbell, Ryan D. Morin

**Author notes:** These authors contributed equally (Krysta M Coyle, Tiana Hillman). Correspondence should be addressed to Dr. Ryan D. Morin and Dr. Krysta M. Coyle., Simon Fraser University, 8888 University Drive, Burnaby BC. V5A 1S6 Canada.

## Abstract

Animal models of human cancers are an important tool for the development and preclinical evaluation of therapeutics. Canine B-cell lymphoma (cBCL) is an appealing model for human mature B-cell neoplasms due to the high sequence similarity in cancer genes to humans and inactive telomerase in adult tissues. We performed targeted sequencing on 86 canine patients from the Canine Comparative Oncology Genomic Consortium, with 61 confirmed as B-cell lymphomas. We confirmed a high frequency of mutations in TRAF3 (45%) and FBXW7 (20%) as has been reported by our group and others. We also note a higher frequency of DDX3X (20%) and MYC (13%) mutations in our canine cohort.

We compared the pattern and incidence of mutations in cBCL to human diffuse large B-cell lymphoma (hDLBCL) and human Burkitt lymphoma (hBL). Canine MYC mutations displayed a focal pattern with 80% of mutations affecting the conserved phosphodegron sequence in MYC box 1, which are known to stabilize MYC protein. We also note that MYC and FBXW7 mutations do not co-occur in our cBCL cohort, leading to the hypothesis that these mutations represent alternative approaches to stabilize MYC in canine lymphoma.

We observed striking differences in the pattern of DDX3X mutations in canine lymphoma as compared to hBL and uncovered a sex-specific pattern of DDX3X mutations in hBL that is not consistent with those identified in canine lymphomas.

In sum, we describe key differences between cBCL and human mature B-cell lymphomas which may indicate differences in the biology of these cancers. This should be considered in future studies of cBCL as a model of human lymphomas.

---

Animal models of human cancers are an important tool for the development and preclinical evaluation of therapeutics. Canine B-cell lymphoma (cBCL) is an appealing alternative to murine preclinical models due to its frequent, spontaneous incidence and clinical and histological similarities to some human mature B-cell neoplasms.^1,2^ Dogs are particularly relevant for comparative oncology as they show a higher sequence similarity in cancer genes to humans relative to mice and telomerase is largely inactive in adult dog tissues, as in humans.^3,4^ Current veterinary care for cBCL includes many of the same chemotherapeutic agents used for human B-cell lymphomas, and the accelerated lifespan of dogs and relative acceleration in cancer progression may allow more rapid observations of experimental treatments.^5–7^

The most common form of cBCL resembles human diffuse large B-cell lymphoma (hDLBCL)^3^ with other subtypes, including Burkitt-like cBCL, less frequently diagnosed.^8,9^ Genomic characterization of hDLBCL continues to reveal novel subtypes with different clinical features and responses to therapy.^10^ Given the mutation patterns that underlie molecular heterogeneity in hDLBCL, we hypothesized that the molecular heterogeneity of cBCL and its relationship to hDLBCL remains incomplete and is not adequately captured by current diagnostic methods.^11^ Moreover, the utility of cBCL as a veterinary model of human disease would be bolstered by an enhanced understanding of the genetic alterations that collectively underlie cBCL.

We obtained fresh frozen and matched plasma/serum from 86 patients from the Canine Comparative Oncology Genomic Consortium (CCOGC), with 61 confirmed as B-cell lymphoma by immunophenotyping (Supplemental Table 1). We extracted total RNA and DNA from 29 tumor samples and performed RNA-seq as previously described.^12^ Genomic DNA was extracted from the remaining tumors using either the AllPrep DNA/RNA Universal Kit or the DNeasy Blood and Tissue Kit (Qiagen). DNA was extracted from plasma or serum using the MagMAX Cell-free DNA Isolation Kit (Thermo Fisher, Waltham, MA).

We aligned RNA-seq reads to the canFam3 reference using STAR^13^ and identified SNVs and indels as described previously.^12^ After identifying genes with evidence for recurrent mutations, we performed targeted sequencing of candidate mutations using custom PCR primers. We produced custom capture baits by PCR amplification of each exon of interest using genomic DNA from a healthy dog as a template.^14^ Tumor DNA was prepared into libraries using the QIAseq FX DNA Library Kit (Qiagen). Plasma and serum DNA was prepared into libraries using the NebNext Ultra II DNA Library Prep Kit (New England BioLabs) followed by enrichment using our baits. We aligned reads to canFam3.1 and visually confirmed mutations using Geneious. Variants were annotated with Variant Effect Predictor and human-dog pairwise alignments were extracted from Ensembl to identify human positions for all canine variants.

Using mutations identified in either tumor or circulating tumor DNA (ctDNA) (Figure S1), we confirmed 9 recurrently mutated genes in canine B-cell lymphoma (Figure 1A), including the previously noted high frequency of mutations in *TRAF3* (45%) and *FBXW7* (20%; Figure 1A).^12,15,16^ *DDX3X* (20%) and *MYC* (13%) were mutated at a higher frequency than has been previously described in cBCL.^15^ These higher rates can be attributed, in part, to our prior observation of high levels of tumor DNA contaminating some of the normal samples.^12^

**Figure 1:**
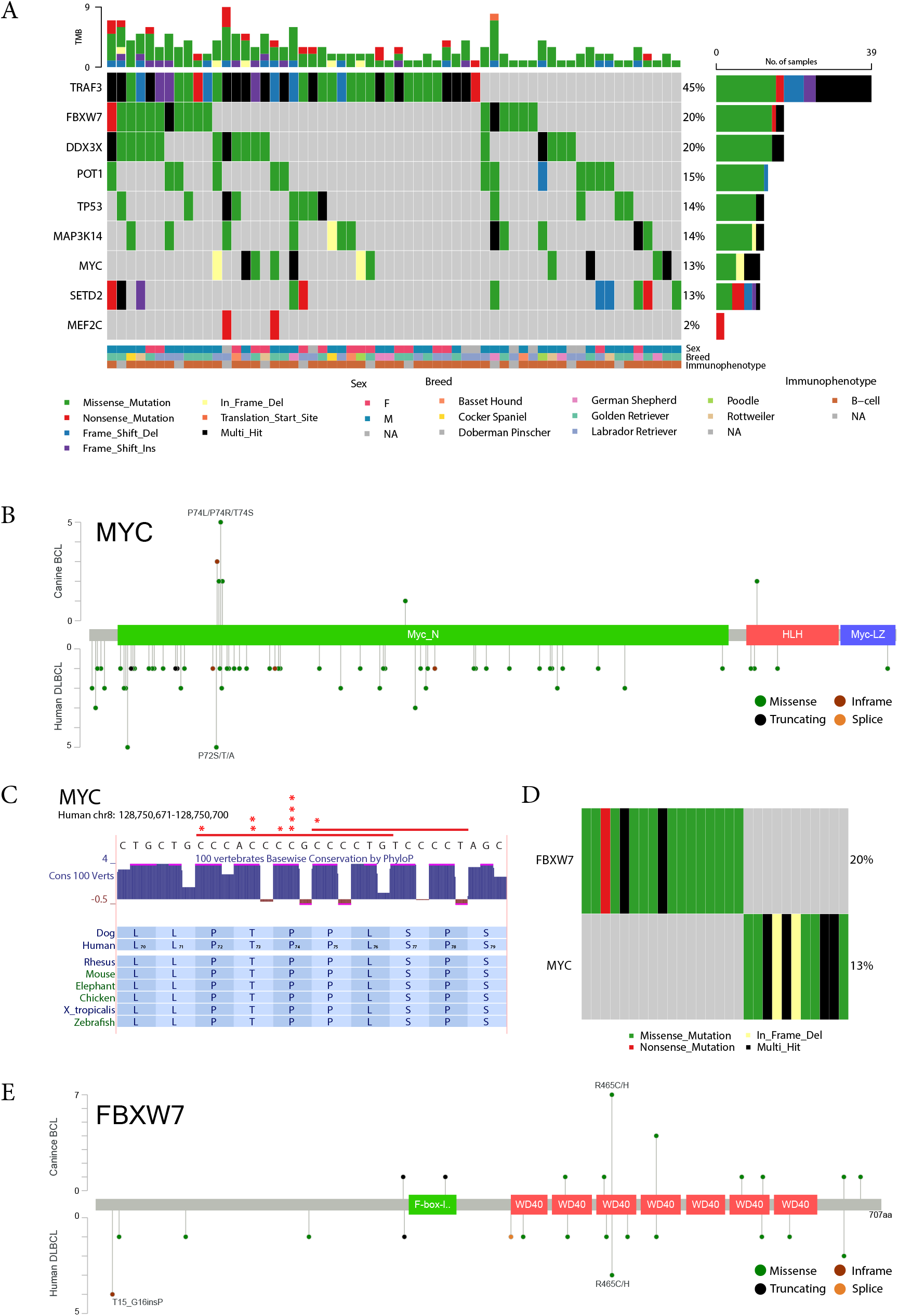
Targeted sequencing of cBCL identifies frequent mutations affecting MYC stability. A. Frequently mutated genes identified in cBCL. Mutations observed across 86 canine BCL samples in 9 genes. After removing suspected germline variants, cBCLs had between zero and nine mutations (mean 2.02) in genes of interest. Mutation frequencies of *POT1* (15%), *TP53* (14%), and *SETD2* (13%) are similar to those reported in previous studies. ^15,16^ *MAP3K14* mutations occur in 14% of cases; however, its frequency in other studies has not been reported.^15^ B. Spatial distribution of mutations observed in *MYC*, compared to human DLBCL. The odds ratio corresponding to the proportion of *MYC* hotspot mutations in cBCL versus hDLBCL is 30.79 (95% CI: 6.73-202.3, p = 1.21 × 10^−7^). C. The *MYC* phosphodegron sequence is highly conserved in vertebrates and the most common site of *MYC* mutations in cBCL (12/15 mutations). D. *MYC* and *FBXW7* mutations do not co-occur in cBCL. E. Spatial distribution of mutations observed in *FBXW7*, as compared to human DLBCL. The hotspot (present in both human and dog BCL) occurs in a WD40 repeat, which forms one of the blades of the beta-propellor and affect a residue forming part of the substrate recognition domain.

We compared the pattern and incidence of mutations between cBCL, hDLBCL, and human Burkitt lymphoma (hBL), from a variety of in-house and published sources (Figure S2).^17–19^ *MYC* is commonly deregulated by translocation in hDLBCL and hBL and these events are commonly associated with point mutations due to aberrant somatic hypermutation.^20^ We observed a low frequency of *MYC* mutations in our cBCL cohort with a more focal pattern that is not consistent with the pattern in hDLBCL (Figure 1B). Twelve mutations (80%) affect the MYC box I, located in a conserved Cdc4 phosphodegron (CPD) sequence (Figure 1C). In human MYC, these are known to stabilize the protein by rendering it resistant to FBXW7-mediated degradation.^21,22^

*FBXW7* mutations are of particular interest as we never observe both *MYC* and *FBXW7* mutations in cBCL (Figure 1D). The most recurrent *FBXW7* mutation affected R470, corresponding to the human R465 codon, which is also a hot spot in hDLBCL (Figure 1E). These mutations are predicted to yield a dominant negative form of FBXW7 that is unable to effectively degrade target substrates including MYC and NOTCH1.^23^ We hypothesize that mutations in *FBXW7* and the *MYC* CPD represent alternative approaches to stabilize MYC, possibly resulting in overexpression.

*DDX3X* was one of the most frequently mutated genes in our cohort (20%) and is among the most frequently mutated genes in hBL (46%) but with a strikingly different pattern (Figure S2). Interestingly, only missense mutations are observed in cBCL whereas hBLs include a high proportion of truncating mutations (Figure 2A).^24^ We explored various clinical features of hBL patients to identify possible explanations for this difference. Separating *DDX3X* mutations from male and female hBLs resolved a similar pattern only in female hBL (Figure 2B) whereas stratification on Epstein-Barr viral status showed no clear pattern (Figure S3). In males, mutations were found across the entire length of the *DDX3X* coding region with a large proportion of truncating mutations including both nonsense and frameshift, whereas the pattern in females is predominantly missense mutations affecting the DEAD box and helicase domains. In contrast, though all *DDX3X* mutations in cBCL are missense, there is no sex bias observed in frequency or location (Figure 2C).

**Figure 2:**
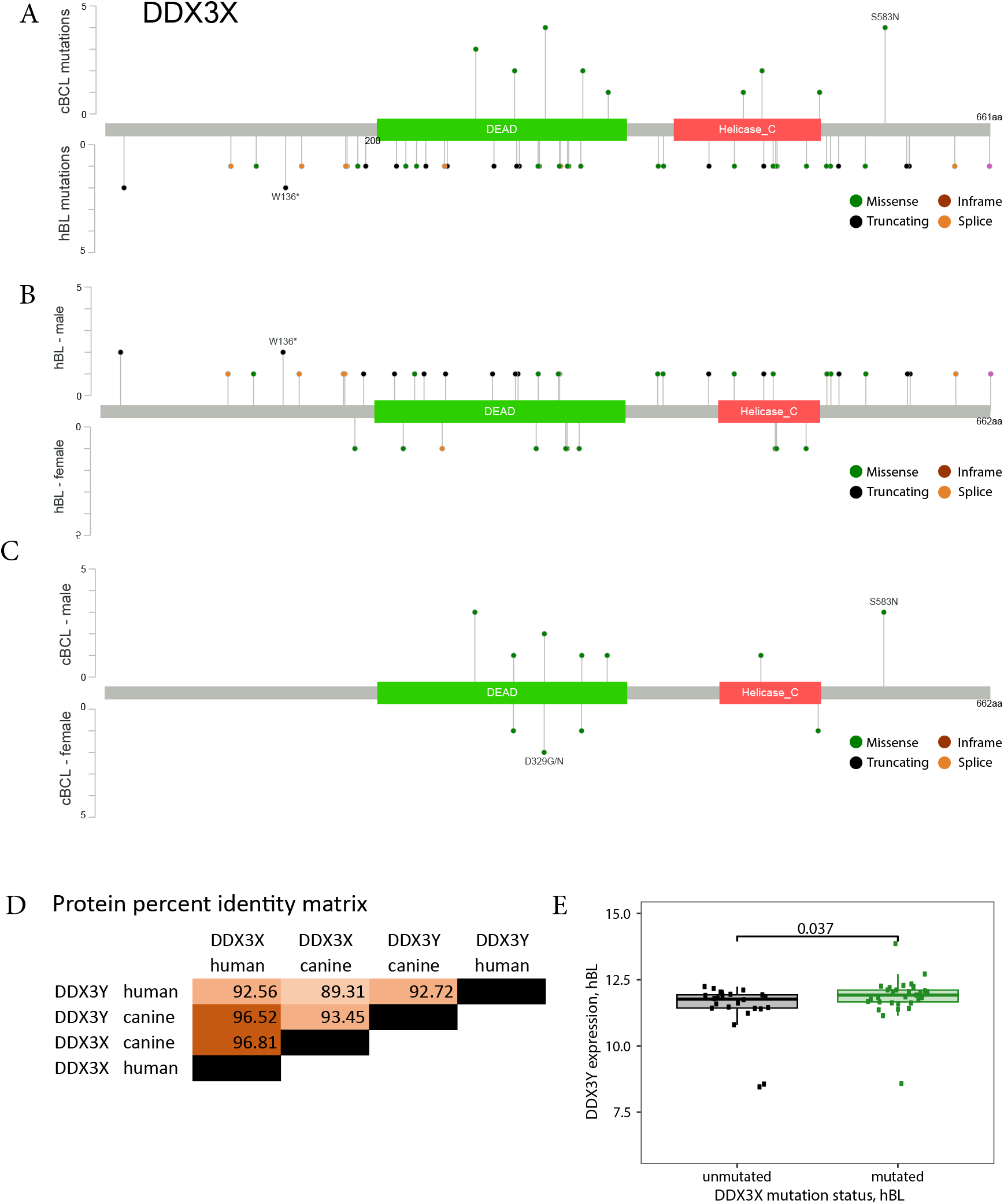
Sex-specific pattern of DDX3X mutations in hBL is not replicated in cBCL. A. Spatial distribution of mutations observed in *DDX3X*, compared to human BL. B. Spatial distribution of *DDX3X* mutations in human male and human female BL. The odds ratio corresponding to the presence of truncating mutations in male BL is infinite (95% CI: 1.44-Inf, p = 0.00918). C. Spatial distribution of DDX3X mutations in canine male and canine female BCL. D. Protein percent identity, calculated by Clustal Omega, is highly similar between human and canine *DDX3X* and the Y-linked paralog *DDX3Y*. E. Expression of *DDX3Y* mRNA is significantly higher in human male BL when a mutation in *DDX3X* is present.

A paralog of *DDX3X*, *DDX3Y*, is encoded on the Y chromosome. Based on high sequence similarity (Figure 2D) and functional evidence, DDX3X and DDX3Y proteins may have partially redundant functions in humans.^24,25^ We considered the possibility that DDX3Y may play a compensatory role in males with *DDX3X* mutations. We found a significantly higher expression of DDX3Y in males hBLs with *DDX3X* mutations when compared to males without these mutations (Figure 2E). A similar comparison was not possible for cBCL due to the small sample size; however, our findings support the premise that these two proteins may have functional redundancy in the context of human lymphomagenesis but may not in canine lymphomagenesis. This represents an important difference between cBCL and human B-cell lymphomas.

cBCL has value as an intermediate between rodent models and clinical trials; however, our data identifies two key factors, namely *FBXW7* and *DDX3X*, that may impact the use of cBCL as a preclinical model for human mature B-cell lymphomas. In human cancers, *FBXW7* is most commonly mutated in cholangiocarcinoma and T-cell acute lymphoblastic leukemia,^23^ but is rarely observed in the mature B-cell malignancies used in this study. We hypothesize that *FBXW7* mutations in cBCL have a redundant function to the mutations affecting the MYC phosphodegron, which may be the cause of the apparent mutual exclusivity observed in this study. This redundancy should be considered in future studies of potential MYC-targeted therapies for canine lymphomas. We also describe a sex-specific pattern of mutations affecting *DDX3X* in hBL which is not recapitulated in cBCL. The discrepancy in mutation patterns between canine and human patients represents an important distinction that may indicate differences in the biology of these cancers

This study has revealed key differences in the mutational profiles of canine and human B-cell lymphomas and provides an impetus for enhanced genomic characterization of canine lymphomas as a model for human NHL, particularly in clinical trial settings.

## Supplemental information attached

## Acknowledgements

This work was supported by an NSERC discovery grant awarded to RDM, a Genome Canada contract, and start-up funds provided by the BC Cancer Foundation to RDM. RDM is a Michael Smith Scholar. KMC is supported by a CIHR Postdoctoral Fellowship. We thank the Canine Comparative Oncology Genomics Consortium (CCOCG) for providing the canine tissue samples. We thank Jovanveer Shoker for his technical contributions to this work.

The Genomic Variation in Diffuse Large B Cell Lymphomas study was supported by the Intramural Research Program of the National Cancer Institute, National Institutes of Health, Department of Health and Human Services. The datasets have been accessed through the NIH database for Genotypes and Phenotypes (dbGaP, accession # phs001444).). A full list of acknowledgements can be found in the supplementary note (Schmitz et al., PMID: 29641966). This work is conducted as part of the Slim Initiative for Genomic Medicine (SIGMA), a joint U.S.-Mexico project funded by the Carlos Slim Health Institute. Data is available at dbGAP (accession # phs000450). The results published here are in whole or part based upon data generated by the Cancer Genome Characterization Initiative (phs000235, Non-Hodgkin Lymphoma project), developed by the NCI. The data used for this analysis are available at https://www.ncbi.nlm.nih.gov/projects/gap/cgi-bin/study.cgi?study_id=phs000235.v6.p1. Information about CGCI projects can be found at https://ocg.cancer.gov/programs/cgci.

## Author Contributions

KMC and RDM prepared the manuscript. KMC, KRB, BMG, and RDM conceived and designed experiments. TH, KRB, MC, SEA, NT, KD, MA, KDL, SW, and KC performed experiments. KMC, TH, MC, BMG, KRB, NT, KD, and RDM performed the analysis and visualization of data.

## References

1. Hansen K, Khanna C. Spontaneous and genetically engineered animal models: Use in preclinical cancer drug development. European Journal of Cancer. 2004;40(6):858–880.

2. Modiano JF, Breen M, Burnett RC, et al. Distinct B-cell and T-cell lymphoproliferative disease prevalence among dog breeds indicates heritable risk. Cancer research. 2005;65(13):5654–61.

3. Rowell JL, McCarthy DO, Alvarez CE. Dog models of naturally occurring cancer. Trends in Molecular Medicine. 2011;17(7):380–388.

4. Nasir L, Devlin P, Mckevitt T, Rutteman G, Argyle DJ. Telomere lengths and telomerase activity in dog tissues: A potential model system to study human telomere and telomerase biology. Neoplasia. 2001;3(4):351–359.

5. Kimmelman J, Nalbantoglu J. Faithful companions: a proposal for neurooncology trials in pet dogs. Cancer Res. 2007;67(10):4541–4544.

6. Hansen K, Khanna C. Spontaneous and genetically engineered animal models; use in preclinical cancer drug development. Eur J Cancer. 2004;40(6):858–880.

7. Honigberg LA, Smith AM, Sirisawad M, et al. The Bruton tyrosine kinase inhibitor PCI-32765 blocks B-cell activation and is efficacious in models of autoimmune disease and B-cell malignancy. Proc Natl Acad Sci U S A. 2010;107(29):13075–13080.

8. Zandvliet M. Canine lymphoma: a review. Veterinary Quarterly. 2016;36(2):76–104.

9. Aresu L, Agnoli C, Nicoletti A, et al. Phenotypical Characterization and Clinical Outcome of Canine Burkitt-Like Lymphoma. Front. Vet. Sci. 2021;8:.

10. Morin RD, Arthur SE, Hodson DJ. Molecular profiling in diffuse large B-cell lymphoma: why so many types of subtypes? Br J Haematol. 2021;

11. Valli VE, Myint MS, Barthel A, et al. Classification of Canine Malignant Lymphomas According to the World Health Organization Criteria. Vet Pathol. 2011;48(1):198–211.

12. Bushell KR, Kim Y, Chan FC, et al. Genetic inactivation of TRAF3 in canine and human B-cell lymphoma. Blood. 2015;125(6):999–1005.

13. Dobin A, Davis CA, Schlesinger F, et al. STAR: ultrafast universal RNA-seq aligner. Bioinformatics. 2013;29(1):15–21.

14. Alcaide M, Rushton C, Morin RD. Ultrasensitive Detection of Circulating Tumor DNA in Lymphoma via Targeted Hybridization Capture and Deep Sequencing of Barcoded Libraries. Methods Mol Biol. 2019;1956:383–435.

15. Elvers I, Turner-Maier J, Swofford R, et al. Exome sequencing of lymphomas from three dog breeds reveals somatic mutation patterns reflecting genetic background. Genome Res. 2015;25(11):1634–1645.

16. Smith PAD, Waugh EM, Crichton C, Jarrett RF, Morris JS. The prevalence and characterisation of TRAF3 and POT1 mutations in canine B-cell lymphoma. The Veterinary Journal. 2020;266:105575.

17. Schmitz R, Wright GW, Huang DW, et al. Genetics and Pathogenesis of Diffuse Large B-Cell Lymphoma. N Engl J Med. 2018;378(15):1396–1407.

18. Chapuy B, Stewart C, Dunford AJ, et al. Molecular subtypes of diffuse large B cell lymphoma are associated with distinct pathogenic mechanisms and outcomes. Nature Medicine. 2018;24(5):679–690.

19. Grande BM, Gerhard DS, Jiang A, et al. Genome-wide discovery of somatic coding and noncoding mutations in pediatric endemic and sporadic Burkitt lymphoma. Blood. 2019;133(12):1313–1324.

20. Khodabakhshi AH, Morin RD, Fejes AP, et al. Recurrent targets of aberrant somatic hypermutation in lymphoma. Oncotarget. 2012;3(11):1308–1319.

21. Welcker M, Orian A, Jin J, et al. The Fbw7 tumor suppressor regulates glycogen synthase kinase 3 phosphorylation-dependent c-Myc protein degradation. Proceedings of the National Academy of Sciences of the United States of America. 2004;101(24):9085–90.

22. Yada M, Hatakeyama S, Kamura T, et al. Phosphorylation-dependent degradation of c-Myc is mediated by the F-box protein Fbw7. The EMBO Journal. 2004;18(3):717–726.

23. Akhoondi S, Sun D, Lehr N von der, et al. FBXW7/hCDC4 Is a General Tumor Suppressor in Human Cancer. Cancer Res. 2007;67(19):9006–9012.

24. Gong C, Krupka JA, Gao J, et al. Sequential inverse dysregulation of the RNA helicases DDX3X and DDX3Y facilitates MYC-driven lymphomagenesis. Molecular Cell. 2021;

25. Venkataramanan S, Gadek M, Calviello L, Wilkins K, Floor S. DDX3X and DDX3Y are redundant in protein synthesis. RNA. 2021;rna.078926.121.

