## Supplemental materials for "Shared and distinct genetic features in human and canine B-cell lymphomas"

Supplemental Tables and Figures

**Table S1. Clinical features of canine cohort.**

| | | N | % of cohort |
|---|---|---|---|
| **Total** | | 86 | 100% |
| **Sex** | | | |
| | M | 47 | 54.7% |
| | F | 26 | 30.2% |
| | unk. | 13 | 15.1% |
| **Immunophenotype** | | | |
| | B-cell | 61 | 70.9% |
| | unk. | 25 | 29.1% |
| **Breed** | | | |
| | Golden Retriever | 31 | 36.0% |
| | Labrador Retriever | 28 | 32.6% |
| | German Shepherd | 8 | 9.3% |
| | Basset Hound | 5 | 5.8% |
| | Poodle | 5 | 5.8% |
| | Rottweiler | 4 | 4.7% |
| | Cocker Spaniel | 2 | 2.3% |
| | Doberman Pinscher | 1 | 1.2% |
| | Unknown | 2 | 2.3% |

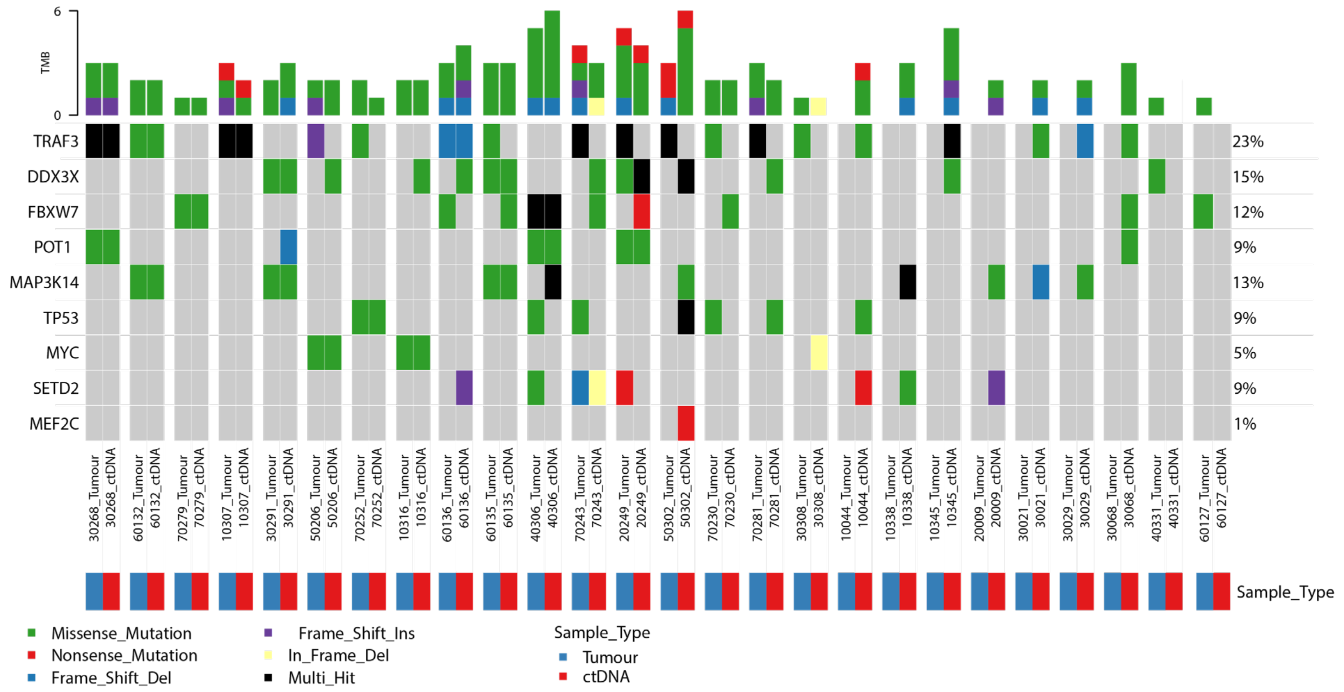


Figure S1. Comparison of mutations between matched tumor and ctDNA samples from canine patients.

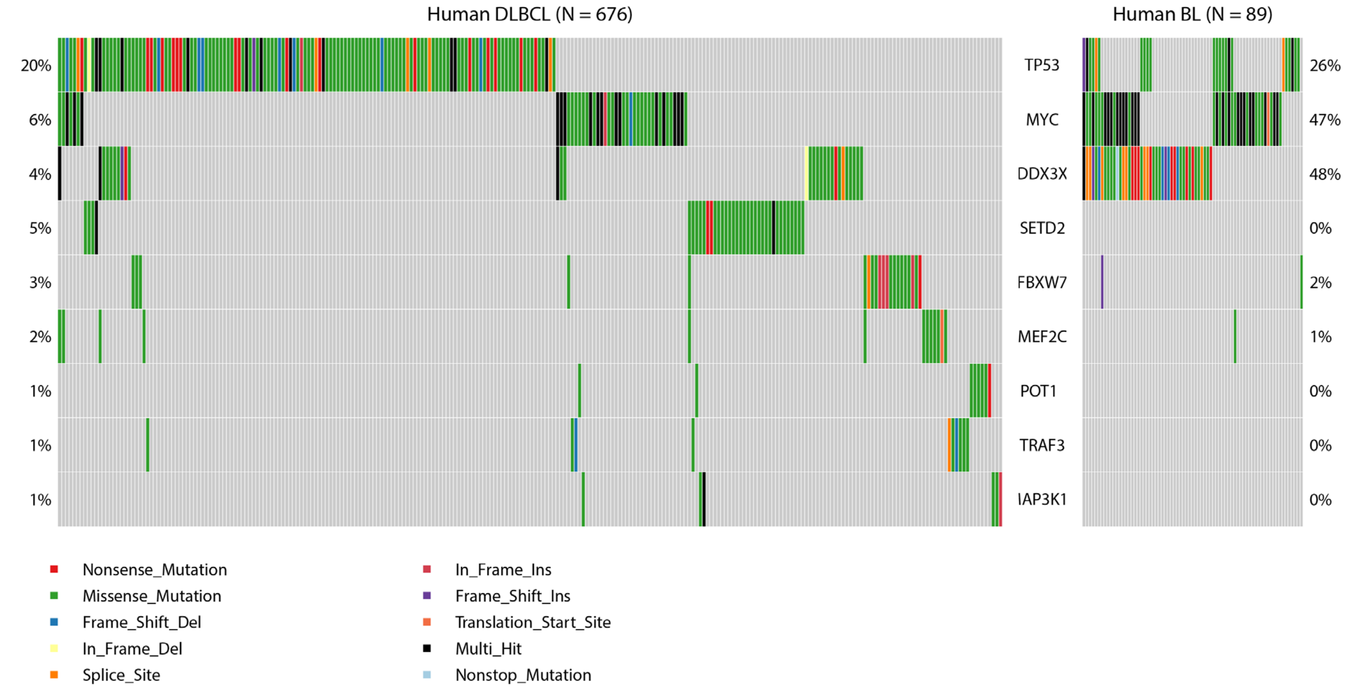


Figure S2. Comparison of mutation frequency in human Burkitt lymphoma (hBL) and human diffuse large B-cell lymphoma (DLBCL). For 30 patients, we sequenced both plasma and tumor samples to determine the consistency between circulating tumor DNA and tumor tissue. Due to the variable discordance between the mutations identified in plasma and tumor samples from the same patient, we show the union of all mutations found in all samples from each patient.

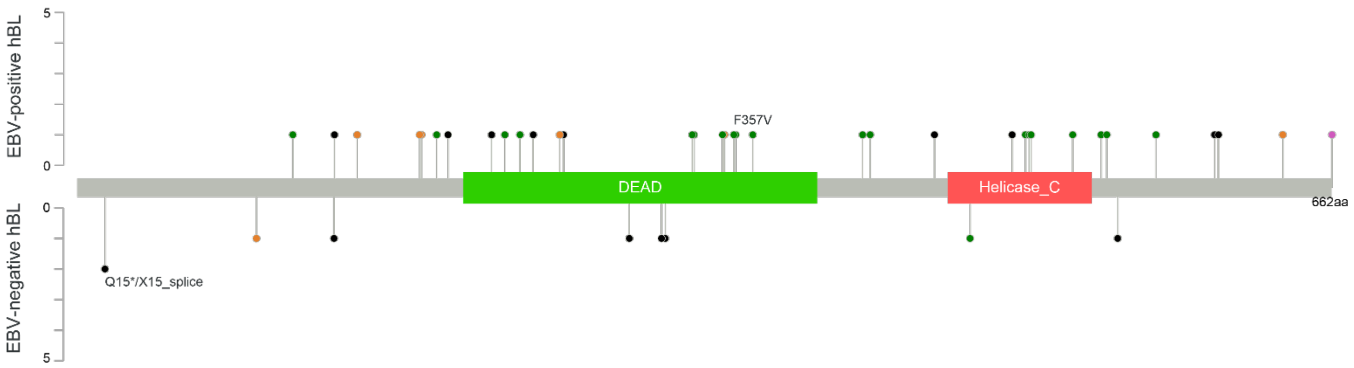


Figure S3: Spatial distribution of *DDX3X* mutations in hBL, separated by EBV status. Truncating mutations (black) are present in both EBV-positive and EBV-negative hBL.

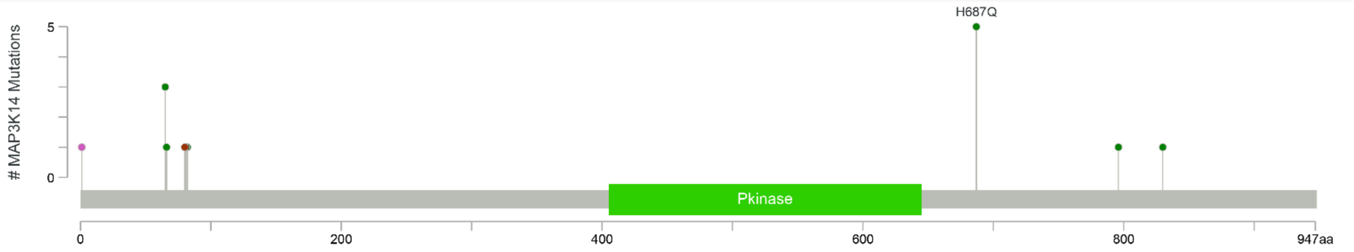


Figure S4: Spatial distribution of *MAP3K14* mutations in cBCL. Mutations are plotted relative to human MAP3K14 amino acids.

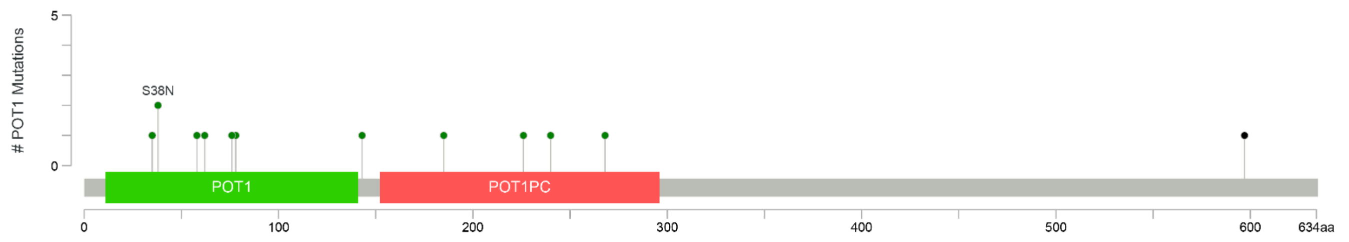


Figure S5: Spatial distribution of *POT1* mutations in cBCL. Mutations are plotted relative to human POT1 amino acids.

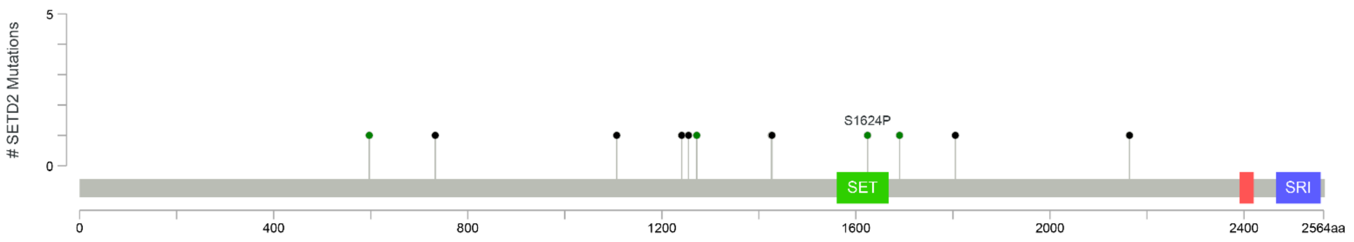


Figure S6: Spatial distribution of *SETD2* mutations in cBCL. Mutations are plotted relative to human SETD2 amino acids.

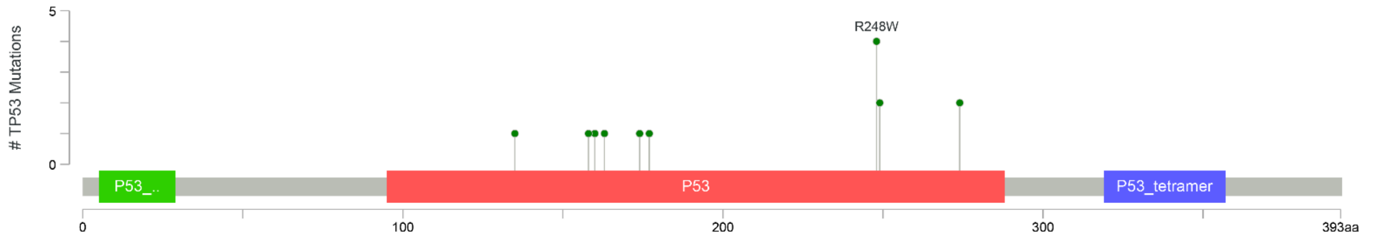


Figure S7: Spatial distribution of *TP53* mutations in cBCL. Mutations are plotted relative to human TP53 amino acids.

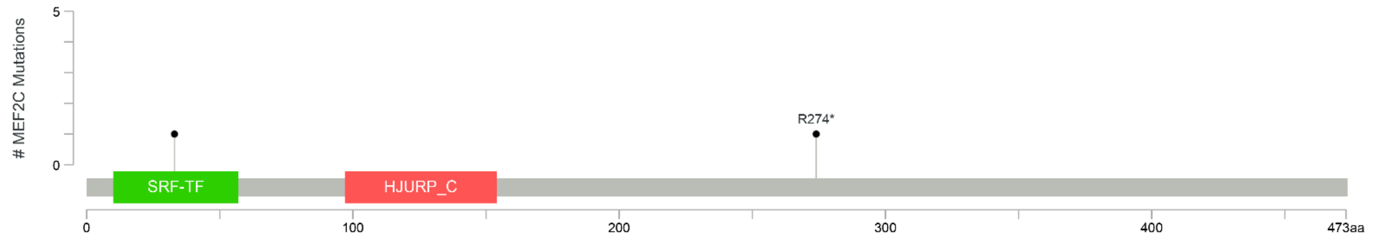


Figure S8: Spatial distribution of *MEF2C* mutations in cBCL. Mutations are plotted relative to human MEF2C amino acids.
